# A widespread decrease of chromatin accessibility in age-related macular degeneration

**DOI:** 10.1101/264564

**Authors:** Jie Wang, Cristina Zibetti, Peng Shang, Srinivasa R. Sripathi, Pingwu Zhang, Marisol Cano, Thanh Hoang, Shuli Xia, Hongkai Ji, Shannath L. Merbs, Donald J. Zack, James T. Handa, Debasish Sinha, Seth Blackshaw, Jiang Qian

## Abstract

Age-related macular degeneration (AMD) is a leading cause of blindness in the elderly. The extent to which epigenetic changes regulate AMD progression is unclear. Here we globally profiled chromatin accessibility in the retina and retinal pigmented epithelium (RPE) from AMD patients and controls. Global decreases in chromatin accessibility occurr in RPE in early AMD, and in the retina with advanced disease, suggesting that dysfunction in RPE cells drives disease progression. Footprints of photoreceptor and RPE-specific transcription factors are enriched in differentially accessible regions (DARs). Genes associated with DARs show altered expression in AMD. Cigarette smoke treatment of RPE cells recapitulates epigenomic changes seen in AMD, providing an epigenetic link between the known risk factors for AMD and AMD pathology. Finally, overexpression of *HDAC11* is partially responsible for the reduction in chromatin accessibility, identifying potential new targets for treatment of AMD.

---

Age-related macular degeneration (AMD) is by far the most common cause of irreversible visual impairment in people over 60^1^. The estimated number of people with AMD in 2020 is 196 million, and will increase substantially with the aging of the global population^2^. The disease is characterized by the early appearance of drusen, pigmentary abnormalities of the retinal pigment epithelium (RPE), and progressive photoreceptor dysfunction that is restricted primarily in the macula, a 6 mm diameter region of the fundus^3^. Although treatments aimed at inhibiting blood vessel growth can effectively slow the progression of the “wet” AMD, no useful treatments exist for the atrophic (“dry”) form of the disease, which account for 90% of all AMD cases^4^.

Currently, GWAS analysis has identified at least 34 AMD genetic risk loci involved in the regulation of complement pathway and inflammation^5,6^. However, these genetic variants can only explain a subset of AMD cases, suggesting a substantial role of environmental factors in the pathogenesis of AMD. Indeed, studies have linked variables such as cigarette smoking and obesity to AMD susceptibility, both of which are known to induce cellular stress and inflammation in a wide range of tissues^7,8^. However, no comprehensive studies have yet been reported on global epigenetic changes associated with AMD progression. This in part, reflects the lack of widely-accepted animal models for AMD^4^, as well as the difficulty in obtaining sufficient amounts of human pathological tissue for analysis. Here we observed global and progressive decreases in chromatin accessibility associated with AMD progression. Both cigarrette smoking treatment and overexpression of the epigenetic regulator *HDAC11* in human iPSC-derived RPE recapitulated the changes in chromatin accessibility. These findings suggest that global decreases in chromatin accessibility may play a critical role in the onset and progression of AMD.

## Landscape of chromatin accessibility in the retina and RPE

In this study, we obtained 16 eyes from 5 controls and 5 AMD donors (Extended Data Table 1). We collected retina and RPE tissues from the macular and peripheral regions of each donor eye, which altogether yielded a total of 19 normal, 9 early AMD, and 17 late AMD-derived samples (Table 1). Although the procurement time is slightly longer for normal samples, major characteristics including gender and age are comparable among normal and AMD samples. Disease status was confirmed with visual examination by an expert observer (J.T.H.).

**Table 1.** The characteristics of ATAC-Seq samples

| Variable | Tissue | Normal | Early AMD | Late AMD | P value* |
| --- | --- | --- | --- | --- | --- |
| No. of samples | Retina | 11 (58%) | 5 (56%) | 9 (53%) | 0.99 |
|  | RPE | 8 (42%) | 4 (44%) | 8 (47%) | 0.99 |
| Region (macula) | Retina | 5 (45%) | 2 (40%) | 5 (56%) | 0.99 |
|  | RPE | 3 (38%) | 2 (50%) | 6 (75%) | 0.38 |
| Gender (male) | Retina | 5 (45%) | 2 (40%) | 2 (22%) | 0.57 |
|  | RPE | 5 (63%) | 1 (25%) | 2 (25%) | 0.41 |
| Age (years) | Retina | 84: 79 ~ 92 | 87: 82 ~ 87 | 90: 90 ~ 94 | 0.09 |
|  | RPE | 88: 84 ~ 92 | 92: 85 ~ 97 | 90: 89 ~ 94 | 0.32 |
| Interval (hours)& | Retina | 10.1 | 5.3 | 6.6 | 0.01 |
|  | RPE | 9.5 | 4.9 | 6.8 | 0.08 |

\*Fisher's exact test and one-way ANOVA were performed respectively. &Interval indicates the time from death to procurement of eye.

To study the global epigenetic landscape of AMD, we used the assay for transposase-accessible chromatin using sequencing (ATAC-Seq) to detect genomic chromatin accessibility, which depicts the active (i.e. open) chromatin and inactive (i.e. condensed) chromatin^9^. We obtained an average of 78.5% mappability and 35.8 million qualified fragments per sample (Extended Data Table 2). ATAC-Seq data from two replicate samples, obtained from adjacent regions in peripheral retina of the same eye, showed high correlation (R = 0.98, Extended Data Fig. 1a), indicating that ATAC-Seq can reliably and reproducibly measure chromatin accessibility in these samples. In total, 78,795 high-confidence open chromatin regions (or peaks) were identified across all retinal samples, and 49,217 peaks were identified across all RPE samples, representing a total of 93,863 distinct peaks. Chromatin accessibility in the retina is overall higher than that of the RPE, potentially reflecting the much greater diversity of cell types in the retina relative to the RPE (Fig. 1a). Comparison of samples from the macular vs. peripheral retina, as well as the macular vs. peripheral RPE, showed broadly similar profiles of chromatin accessibility (Fig. 1a).

**Figure 1.**
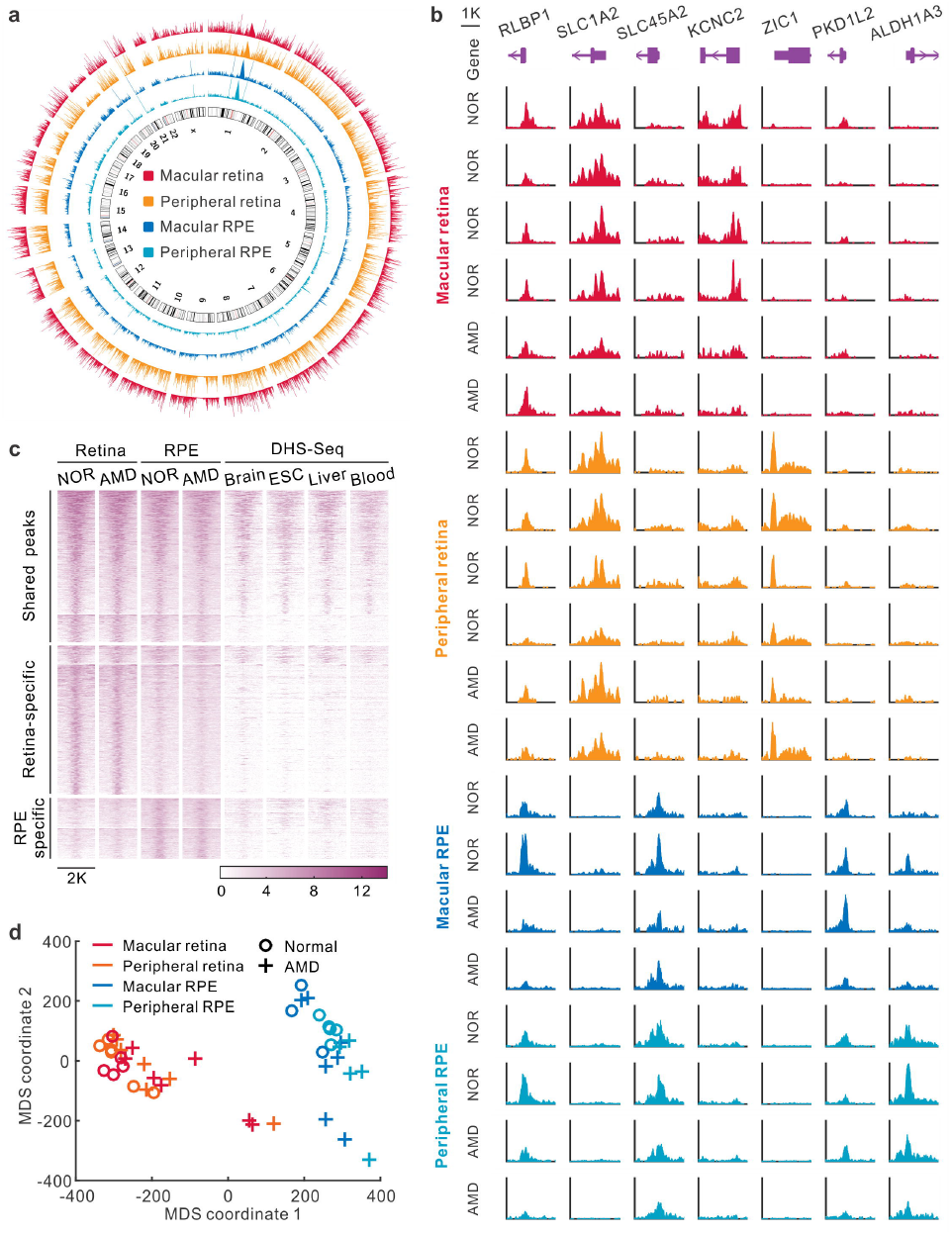
The landscape of chromatin accessibility in human retina and RPE. (a) Genome-wide chromatin accessibility of a control eye. (b) The instances of open chromatin in the retina and RPE from healthy controls (NOR) and AMD patients. (c) Specific and shared ATAC-Seq peaks in the retina and RPE. Each row represents one peak. The color represents the intensity of chromatin accessibility. Peaks are grouped based on K-mean clustering and aligned at the center of regions. (d) Multidimensional scaling (MDS) of all retina and RPE samples.

The data revealed categories of peaks that are either specific to, or shared between, the retina and RPE. For instance, a peak associated with *RLBP1* is shared by the retina and RPE, whereas a peak associated with *SLC1A2* is specific to the retina, and another peak within the *SLC45A2* gene is RPE-specific (Fig. 1b). Furthermore, a small number of region-specific peaks (e.g. peaks in *KCNC2* and *ZIC1*) are selectively detected in the macular and peripheral retina, respectively, while others (e.g. peaks in *PKD1L2* and *ALDH1A3*) are selectively accessible in macular and peripheral RPE (Fig. 1b). Overall, 39,394 (42.0%) ATAC-Seq peaks are shared by the retina and RPE, 38,625 (41.1%) peaks are retina-specific, and 15,844 (16.9%) peaks are RPE-specific (Fig. 1c). We identified 5,855 increased and 2,689 decreased peaks in the macular retina, relative to the paired retinal samples from the peripheral region (Extended Data Fig. 1b). Meanwhile, 432 increased and 959 decreased peaks were detected in macular RPE relative to peripheral RPE (Extended Data Fig. 1c). We observed that a great majority (81.7%) of the peaks that are shared between the retina and RPE are also detected in other tissues (Fig. 1c). In contrast, only 4,626 (12%) of retina-specific peaks and 7,644 (48.2%) of RPE-specific peaks are detected in other tissues, implying that these peaks potentially represent highly tissue-specific *cis*-regulatory elements.

We then calculated the overall similarity of the ATAC-Seq profiles among all the samples using multidimensional scaling. As expected, this analysis showed that the samples are clustered into two groups, one from the retina and the other from the RPE (Fig. 1d). Moreover, most AMD samples are clearly separated from normal samples, suggesting an extensive difference in chromatin accessibility between healthy and AMD tissues.

## Chromatin accessibility is broadly decreased in the retina and RPE of AMD samples

To explore the impact of AMD, we analyzed the differences in chromatin accessibility between normal and AMD retinas. When comparing the accessibility profiles, we noticed substantial quantitative differences in peak signal between normal and AMD retina samples. For example, in three known regulatory regions of the rhodopsin gene *RHO*, chromatin accessibility is progressively decreased from normal to early-stage, and then to late-stage AMD (P < 0.05, Fig. 2a). By comparing the signal for each peak in healthy and AMD samples from both macular and peripheral retinas, we observed that 72,689 (92.3%) peaks have reduced chromatin accessibility in AMD (Fig. 2b). These quantitative differences in chromatin accessibility do not result from the process of normalizing ATAC-Seq data because different normalization approaches gave similar results (Extended Data Fig. 1d). Moreover, we separated retina samples into two groups from the macular and peripheral regions. Relative to the peripheral region, we observed more intense global decrease of chromatin accessibility in the macular region of AMD retina (94.5% for macula and 79.9% for periphery with the reduced chromatin accessibility, Extended Data Fig. 1e and 1f).

**Figure 2.**
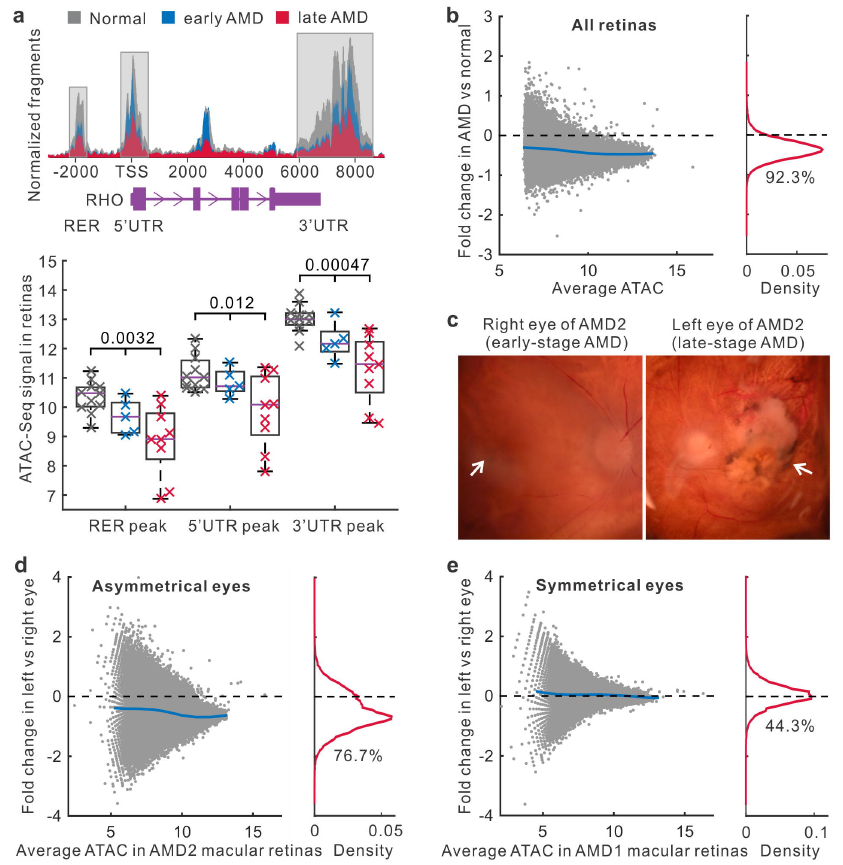
Changes of chromatin accessibility in AMD retinas. (a) The chromatin accessibility in regulatory regions of the rhodopsin gene *RHO*. TSS, transcript start site. RER, rhodopsin enhancer region. UTR, untranslated region. (b) Changes of chromatin accessibility in AMD relative to normal in all retina samples. Each dot represents one ATAC-Seq peak. Blue line indicates average fold changes of peaks. The percentage of reduced peaks is shown under the density curve. (c) The microscopy of right and left eyes from one AMD patient with asymmetrical disease status. The left image shows early AMD (mild RPE pigmentary changes) while the right image shows geographic atrophy. Arrows indicate the macular regions. (d) ATAC-Seq signal changes in right and left macular retinas from the AMD patient with asymmetrical disease status. (e) Accessibility changes in one AMD patient whose eyes are at the same disease stage.

To extend this observation, we obtained a pair of eyes from a donor whose AMD status was asymmetrical, with the right eye showing early-stage AMD, and the left eye showing late-stage dry AMD (Fig. 2c). By comparing these eyes, we excluded the contribution of potential genetic and environmental differences that might complicate the analysis of epigenetic changes associated with AMD progression. Interestingly, a large number (76.7%) of peaks in the macular retina from the more severely affected eye had decreased intensities relative to the less severely affected eye (Fig. 2d). For the donors whose left and right eyes had been diagnosed at the same disease stage, the chromatin accessibility profiles were highly symmetrical in their macular retinas (Fig. 2e and Extended Data Fig. 2a). This analysis confirmed that a widespread decrease in chromatin accessibility is associated with AMD progression.

Next, we analyzed changes of RPE chromatin accessibility in AMD. In all RPE samples, a great number (91.6%) of peaks showed the reduced intensity in AMD relative to normal samples (Fig. 3a). This reduction in intensity associated with AMD RPE was observed in both macular and peripheral regions (88.6% for macula and 94.1% for periphery showed reduced ATAC-Seq signal) (Extended Data Fig. 2b and 2c). In the RPE from the patient who showed different stages of AMD between eyes, the intensities of 42,860 (87.1%) peaks were reduced in the more severely affected left eye (Fig. 3b). In contrast, a symmetrical distribution was observed in donors where both eyes were in the same disease stage (Fig. 3c and Extended Data Fig. 2d). Taken together, our data show a widespread decrease in chromatin accessibility that is observed in both the retina and RPE from AMD patients.

**Figure 3.**
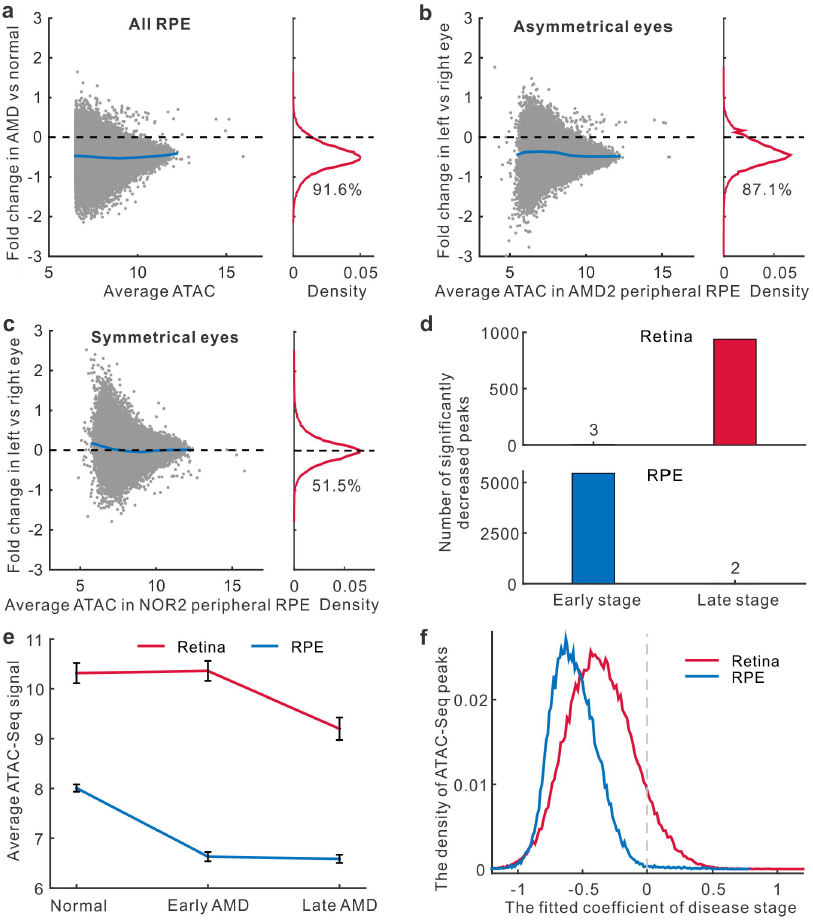
Changes in chromatin accessibility in the RPE and at different disease stages.(a) Changes of chromatin accessibility in AMD RPE for all samples. The percentage of reduced peaks is showed under the density curve. (b-c) Changes of chromatin accessibility in the RPE from donors whose eyes are at the different or the same stage of disease. (d) The number of peaks in the retina and RPE with significantly decreased accessibility for early AMD vs normal (early stage) or late AMD vs early AMD (late stage). (e) Average signal of ATAC-Seq peaks differentially accessible at any stage of AMD. Error bars represent the standard error of mean. (f) The density of the stage coefficients in the fitting model of retina and RPE ATAC-Seq peaks.

## The retina and RPE show decreased chromatin accessibility at different stages of AMD progression

We further set out to identify changes in ATAC-Seq peak intensity that were associated with disease stage in both the retina and RPE. When we compared 5 retinal samples obtained from early-stage AMD to 11 retinal samples obtained from healthy controls, we observed only 3 statistically significant decreases in peak intensity (Fig. 3d and Extended Data Fig. 3a). However, by comparing 9 late-stage AMD retinal samples to these same 5 early-stage retinal samples, we observed 939 peaks with significantly decreased intensity, suggesting that the chromatin accessibility changes in the retina occur only during late stages of disease.

In contrast, when we compared 4 RPE samples from early-stage AMD to RPE samples from 8 healthy controls, we observed 5,458 significantly decreased peaks, but observed only 2 significantly decreased peaks when these same early-stage samples were compared to 8 RPE samples from late-stage AMD (Fig. 3d and Extended Data Fig. 3a). Likewise, when averaging the intensities of significantly decreased peaks at any stage of AMD progression, we found the striking decrease of chromatin accessibility in the RPE occurred at an earlier disease stage than that observed in the retina (Fig. 3e). This observation fits with the widely accepted theory that changes in RPE function trigger AMD^10^, and suggests that epigenetic changes in RPE cells might be a critical factor that regulates disease onset.

## AMD-associated changes in gene regulatory networks

We next sought to determine the functional consequence of the differentially accessible regions (DARs) that were observed in normal and AMD samples. To define statistically significant DARs, we used a linear regression model to take into account the potential effects from other confounding factors such as topographical differences (macula vs. periphery), age, gender, and procurement interval. The model estimated the relative contributions from these factors and the effects of disease stage (normal, early, and late AMD) to variations in peak intensity. Our analysis suggested that the peaks in macular retinas are more likely to be reduced than those in peripheral retinas (Extended Data Fig. 3b). Moreover, a longer procurement interval leads to smaller peaks. Given that the procurement interval of AMD samples is slightly shorter than normal samples (Table 1), it is highly unlikely that an altered procurement interval leads to the decreased peak intensity in AMD samples that is observed in this study. Most importantly, the coefficients for disease stage are significantly negative for a large number of the peaks in the retina (38,520 peaks, 48.9%, FDR < 0.05) and in the RPE (41,168 peaks, 83.7%, Fig. 3f and Extended Data Fig. 3b and 3c), suggesting that late stages of disease are associated with lower peak intensity.

For the retina and RPE, we chose the top 5,000 peaks with significantly negative coefficients of disease stage as DARs (set FDR < 0.01 and ranked by the coefficients, Supplementary Table 1 and 2). We examined the genomic location of these DARs and found that retinal DARs are enriched in intergenic regions (Extended Data Fig. 4a). RPE DARs, in contrast, are enriched in promoters. By checking whether transcription factor (TF) binding was affected in the retina and/or RPE, we observed 22 and 13 TF motifs that are strongly enriched in the retinal and RPE DARs, respectively (Fig. 4a and Extended Data Table 3). For example, the binding motifs of OTX2 and CRX, factors known to play an important role in controlling gene expression in photoreceptors^11,12^, are enriched in AMD retinas. Moreover, OTX2 showed a significantly decreased footprint in DARs for comparison of late-stage AMD to normal samples (Extended Data Fig. 4b).This pattern confirmed that chromatin accessibility of OTX2 target sites is decreased in retinal samples with AMD disease, suggesting that reduced target sites binding by retina and RPE-specific TFs play a critical role in AMD pathogenesis.

**Figure 4.**
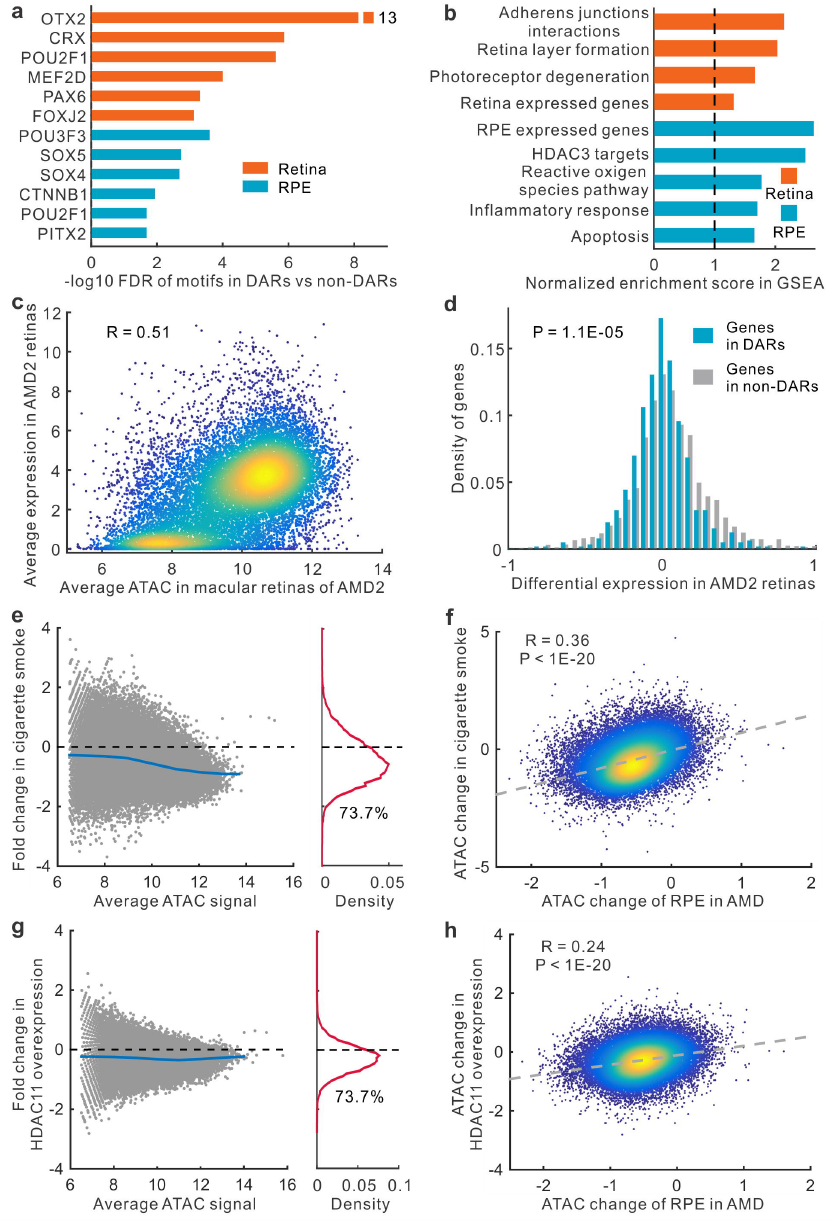
DAR-associated features and chromatin accessibility changes in cigarette smoke-treated or *HDAC11*-overexpressed RPE cells. (a) Enriched TF motifs in footprints within DARs. (b) Significantly enriched functions of DAR nearby genes from gene set enrichment analysis (GSEA). (c) The relationship of chromatin accessibility and RNA-Seq measured gene expression in retinas. (d) The density of differential expression in left vs right retinas of the AMD patient. P value for Student’s t-test is shown. (e) Changes in chromatin accessibility after cigarette smoke treatment. The percentage of reduced peaks is shown under the density curve. (f) Comparison of accessibility changes in AMD RPE and cigarette smoke-treated RPE cells. (g) Changes in chromatin accessibility with *HDAC11* overexpression. (h) Comparison of accessibility changes in AMD RPE and *HDAC11*-overexpressed RPE cells. The grey dashed line is the fitting line. R is Pearson’s correlation coefficient.

## Genes in DARs show altered expression in AMD

We further tested whether the expression levels of genes associated with DARs are more likely to be altered in AMD. We first checked that DAR-associated genes in both the retina and RPE were highly enriched for genes that were selectively expressed in each tissue (Fig. 4b). Moreover, DAR-associated genes in the retina were substantially more likely to regulate retinal layer lamination and photoreceptor survival, while DAR-associated genes in the RPE were more likely to regulate the inflammatory response and apoptosis, which are important biological processes in AMD (Fig. 4b).

Using RNA-Seq data obtained from the patient with differential AMD stages between eyes, we observed that ATAC-Seq peak intensity was highly correlated with gene expression in both the retina and RPE (Fig. 4c and Extended Data Fig. 4c). In the patient with asymmetric AMD progression, DAR-associated genes were significantly more likely to be downregulated in late-stage AMD relative to early-stage AMD (P = 1.1×10^−5^, Fig. 4d and Extended Data Fig. 4d). These results suggest that altered chromatin accessibility in binding sites of retina and RPE-enriched TFs leads to reduced expression of associated genes in AMD.

## Epigenetic changes generally occur independently of AMD risk-associated genetic variants

To examine whether the observed changes in chromatin accessibility resulted from AMD-associated genetic variants, we compared the distribution of DARs to that of genetic variants linked to AMD susceptibility by GWAS analysis^5^. For each DAR, we tested whether it overlapped with one or more AMD-associated SNPs identified by GWAS. Interestingly, we observed that very few of AMD-associated SNPs were covered by DARs in genomic location. There are < 0.1% of all DARs in the retina, and < 0.2% of DARs in the RPE, overlapped with AMD-associated SNPs (Extended Data Fig. 4e). Even if we extend a 5kb window in each direction of DARs, the proportion of DARs overlapped with AMD-associated SNPs was still low (< 0.3% for the retina and < 0.4% for the RPE). For comparison, we also examined the fractions of non-DARs and non-peaks that overlapped with AMD-associated SNPs, and found that the overlap was comparable with those of DAR regions in both retina and RPE. These data imply that the observed differences in chromatin accessibility are unlikely due to local AMD-associated genetic variants.

## Cigarette smoke treatment leads to global decreases of chromatin accessibility in human iPSC-derived RPE cells

Since cigarette smoking is the strongest environmental risk factor for AMD^13^, we tested whether cigarette smoke treatment of cultured human RPE cells could trigger similar changes in chromatin accessibility from AMD samples. The cultured iPSC-derived RPE cells were examined by flow cytometric analysis of RPE-specific markers (Extended Data Fig. 5a). Furthermore, we performed the ATAC-Seq in the cultured RPE cells and confirmed that RPE cells showed a broadly similar pattern of peak distribution to an average ATAC-Seq profile of all healthy samples from RPE tissue (R = 0.83, Extended Data Fig. 5b). For iPSC-derived RPE cells exposed to cigarette smoke extract, ATAC-Seq profile showed a global decrease in chromatin accessibility (Fig. 4e). More importantly, when comparing cigarette smoke treated RPE cells with RPE tissue from AMD patients, we found that the changes in chromatin accessibility are highly correlated (Fig. 4f; R = 0.36, P < 10^−20^). The results indicate that cigarette smoke treatment in RPE cells induces a widespread decrease of chromatin accessibility that is much like that seen in AMD.

## Increased HDAC11 expression leads to global decreases in chromatin accessibility

Next, we attempted to identify the genes that could induce similar change in chromatin accessibility observed during AMD progression. To do so, we queried a previously published collection of microarray data from AMD samples^14^, along with our own RNA-Seq analysis, to identify differentially expressed genes that are known to regulate chromatin accessibility. Among all *HDAC* genes, we found that the class IV histone deacetylase *HDAC11* had significantly increased expression in the RPE during early disease stages (Extended Data Table 4 and Fig. 5c). Furthermore, cigarette smoke treatment of iPSC-derived RPE cells also increased HDAC11 expression based on a western-blot analysis (Extended Data Fig. 5d and 5e).

We then tested whether overexpression of *HDAC11* induces broad decrease in chromatin accessibility. First, we confirmed that *HDAC11* was overexpressed in plasmid transfected RPE cells (Extended Data Fig. 5f). We then examined the chromatin accessibility profile in the monolayers of *HDAC11*-overexpressing RPE cells, and observed a widespread decrease of chromatin accessibility in *HDAC11*-overexpressing cells relative to cells transfected with empty vector (Fig. 4g). Interestingly, the changes induced by *HDAC11* overexpression were strongly consistent with those changes seen in the RPE of AMD patients (Fig. 4h). These results suggest that *HDAC11* overexpression may be partially responsible for the global decreases of chromatin accessibility associated with AMD progression.

## Discussion

In this study, we report the first comprehensive analysis of changes in chromatin accessibility in AMD. These changes in chromatin accessibility are seen first in the RPE, and then later in the retina. Likewise, we observed greater changes in the macular retina than in the peripheral retina. These data fit with the typical pattern of AMD progression, where changes in the RPE precede the dysfunction and death of macular photoreceptors^10^.

Several lines of evidences suggested that the changes in chromatin accessibility are not simply due to cell death in the retina or RPE. First, a diverse proportion of cell type-specific genes (e.g. rods, cones and Müller glia) were associated with decreased ATAC-Seq peaks (Extended Data Fig. 5g), suggesting that the decrease of chromatin accessibility is not simply due to photoreceptor death. Second, the genes associated with DARs are enriched for specific cellular functions, suggesting that the regions for reduced chromatin accessibility are selective. Third, both cigarette smoke treatment and *HDAC11* overexpression mimic the effect of AMD on chromatin accessibility. Altogether, the results suggest that cell death in AMD is unlikely to cause the observed global reduction in chromatin accessibility.

We hypothesize that global changes in chromatin accessibility may be seen in many other diseases. Indeed, large-scale changes in chromatin accessibility have been previously reported in metastatic cancer, where globally increased chromatin accessibility appears to directly drive disease progression^15^. Cancer cells show generally high levels of metabolic activity, and these changes may partially reflect this fact^16,17^. In contrast, neurodegenerative diseases such as AMD are associated with decreased metabolic activity from early stages onward^18,19^. The resulting reduction in cellular levels of acetyl-CoA, an essential cofactor for histone acetylation, that occurs during disease progression may be a common mechanism that contributes to the observed changes in chromatin accessibility ^20^. Our data raise the possibility that global, quantitative reduction in chromatin accessibility may also be observed in other retinal dystrophies, and for neurodegenerative diseases in general.

## Methods

### Human samples

Fresh postmortem eyes were processed within 14 hours after death when obtained from Eye Banks (Portland, USA) and National Disease Research Interchange (Philadelphia, USA). Donor information is summarized in Table S1. The disease conditions were determined by medical record, and the eye globes were further examined by an experienced retinal physician with expertise in AMD (J.T.H.). The retinas were defined as normal when there were no abnormalities observed using a dissecting microscope. Early-stage AMD was defined by the presence of any RPE pigmentary changes and/or large-size drusen (>125μm diameter). Late-stage AMD was defined by areas of geographic atrophy due to loss of the RPE. In this study, we only included dry AMD and excluded wet AMD. For each eye, we separated the retina and RPE and then obtained a punch (6 mm diameter) of retina or RPE tissue in each of macular and peripheral regions. We obtained paired eyes from some donors and one eye from other donors.

### ATAC and RNA sequencing

Biopsy punches of fresh retina and RPE tissues were re-suspended in cold PBS. Chromatin was extracted and processed for Tn5-mediated tagmentation and adapter incorporation, according to the manufacturer’s protocol (Nextera DNA sample preparation kit, Illumina®) at 37 °C for 30 minutes. Reduced-cycle amplification was carried out in presence of compatible indexed sequencing adapters. The quality of the libraries was assessed by DNA-based fluorometric assay (Thermo Fisher Scientific™) and automated capillary electrophoresis (Agilent Technologies, Inc.). Up to 3 samples per lane were pooled and run on a HiSeq2500 Illumina sequencer with a paired-end read of 50bp.

Total cellular RNA was purified using the Qiagen RNAeasy Mini kit and samples with RNA integrity number ≥ 7 were further processed for sequencing. Libraries were prepared using Illumina TruSeq RNA Sample kit (Illumina, San Diego, CA) following manufacturer’s recommended procedure. Briefly, total RNA was denatured at 65°C for 5 minutes, cooled on ice, purified and incubated at 80°C for two minutes. The eluted mRNA was fragmented at 94°C for 8 min and converted to double stranded cDNA, end repaired, A-tailed, and ligated with indexed adaptors and run on a MiSeq Illumina sequencer. The quality of the libraries was also assessed by RNA-based fluorometric assay (Thermo Fisher Scientific™) and automated capillary electrophoresis (Agilent Technologies, Inc.).

### Mapping and normalization of ATAC-Seq

After removing adaptors using Trimmomatic^21^, 50bp paired-end ATAC-Seq reads were aligned to the human reference genome (GRCh37/hg19) using Bowtie2 with default parameters^22^.After filtering reads from mitochondrial chromosome M and sex chromosome Y, we included properly paired reads with high mapping quality (MAPQ score > 10, qualified reads) through SAMTools for further analysis^23^. Duplicate reads was removed using Picard tools MarkDuplicates program (http://broadinstitute.github.io/picard/).

ATAC-Seq peak regions of each sample were called using MACS2 with parameters -- nomodel --shift −100 --extsize 200^24^. The blacklisted regions in human were excluded from peak regions (https://www.encodeproject.org/annotations/ENCSR636HFF/).To get a union set of peaks, we merged the ATAC-Seq peaks of which the distance between proximal ends is less than 10 base pairs. We totally identified 308,019 peaks from retina samples and 208,592 from RPE samples. For each of retina and RPE samples, the fragments were counted across each peak region using HTSeq^25^. We further calculated the normalized fragments (C_N_) by dividing the raw fragments (C_R_) by the library size (S_L_) through the formula: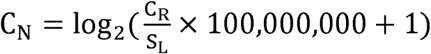. In the study, we used the count of the qualified fragments as the library size for each sample. For downstream analysis, we only included those peak regions with average normalized signal ≥ 6.5 (around the value of 75% quantile) across retina or RPE samples. We obtained 78,795 peaks for retina samples and 49,217 peaks for RPE, consisting of 93,863 distinct peaks. To validate whether different normalization approaches have changed the observations, we alternatively used the count of the properly paired fragments as the library size to normalize fragment counts.

### K-means clustering and multidimensional scaling

The circle plot was performed to visualize genomic ATAC-Seq peaks using Circos^26^. To compare chromatin accessibility of retina and RPE with other tissues, we mapped chromatin accessibility of 125 cells or tissues measured by DHS-Seq to ATAC-Seq peak regions^27^. DHS-Seq data were downloaded from ENCODE project (https://genome.ucsc.edu/ENCODE/index.html). K-means clustering was used to divide ATAC-Seq peaks into tissue-specific and shared groups. To present a two-dimensional distribution of the retina and RPE samples, we performed multidimensional scaling (MDS), in which all pairwise Euclidean distances were calculated as the distance metric. The distances in MDS represent the similarity of samples. The analyses were carried in R platform.

### Comparison of chromatin accessibility and differentially accessible regions (DARs)

An MA plot (log_2_ fold change vs. mean average) was used to visualize the change of chromatin accessibility for all peaks. Meanwhile, the density of peaks was showed across the epigenetic changes. For ATAC-Seq peaks, we accessed the significant change of chromatin accessibility between different groups using edgeR^28^. The count of the qualified fragments in each sample was used as the library size. It was defined as significantly changed if the peak has |log_2_ fold change| > 0.8 and FDR < 0.05. We compared samples in macular region from healthy tissues to the paired samples from peripheral region to identify region-specific ATAC-Seq peaks. For the comparison among three groups of normal, early AMD and late AMD samples, an ANOVA-like test was performed to identify peaks with significant differences (FDR < 0.01 was considered as the significant difference).

In order to estimate the relative contribution of disease stage (normal, early-stage and late-stage) and other confounding factors to the change of ATAC-Seq signal, we conducted linear regression model in which the normalized fragments of ATAC-Seq were taken as the dependent variable. The cofounding factors include the region of tissue (macula and periphery), gender (male and female), age, and procurement interval. Data fitting was performed separately for retina and RPE samples.

The formula for linear regression is following: ATAC-Seq signal = α _0_ + α _1_×stage (0, 1 and 2 separately for normal, early-stage and late-stage) + α _2_×region (−1 and 1 separately for periphery and macula) + α _3_×gender (−1 and 1 separately for female and male) + α _4_×age (years) + α _5_ ×interval(hours)+εIn the formula, the parameter α is the fitted coeffiecient for each variable, which represents the effect of variable on the change of chromatin accessibility. The parameter ε represents random noise. This was performed using the function of linear models ‘lm’ in R platform.

From the peaks with significant association between change of chromatin accessibility and disease stage (using cutoff of FDR < 0.01), we selected top 5,000 peaks with the most negative coefficient of disease stage as differentially accessible regions (DARs) separately for retina and RPE. We excluded DARs and took the rest of ATAC-Seq peaks as non-DARs.

### Genomic features and function enrichment of DARs

We used ANNOVAR to annotate ATAC-Seq peaks with the nearest genes^29^. ATAC-Seq peaks were assigned to four categories, including promoter-proximal, exonic, intronic, 3’UTR and intergenic peaks. Promoter is defined as the region within 2kb of the reference transcript start site (TSS) from UCSC database (https://genome.ucsc.edu/). The peaks located in 5’UTR were also taken as promoter-proximal peaks. The peaks located at downstream 2kb of transcript end site were merged to 3’UTR peaks. Gene-proximal peaks include the peaks within 2kb of the gene body.

Using the linear model, we obtained the coefficient of disease stage for each ATAC-Seq peak and nearby gene. We then ranked genes based on the corresponding coefficient of disease stage. If multiple ATAC-Seq peaks were mapped to the same gene, we assigned the lowest coefficient to the gene. Function enrichment of accessibility change-ranked genes were further performed using GSEA software^30^. Beside of GSEA database, gene sets of photoreceptor degeneration, retina expressed genes, and RPE expressed genes are from published papers^14,31,32^. Similar analysis of GSEA was performed to identify hot spots where are enriched of DAR-associated genes.

### Motifs and footprints of transcription factors associated with DARs

To identify DAR-related transcription factors (TFs), we obtained 1,043 position weight matrices of TF motifs from TRANSFAC database^33^. Considering functional TFs should have gene expression, we filtered TF motifs if gene expression of TF < 5 in both normal and AMD tissues based on microarray data from GSE29801^14^. We thus qualified 485 TF motifs in the retina and 521 motifs in the RPE. We first scanned all potential TF binding motifs (P value < 1.0^-4^) across human genome using FIMO^34^. Meanwhile, we used DNase2TF to identify all potential regulatory regions (the footprints of TFs) in open regions (ATAC-Seq peaks) separately from retina and RPE samples^35^. Within the footprints from DARs and non-DARs, we separately identified the number of TF motifs. Hypergeometric test was further performed to assess the enrichment of TF motifs in the footprints of DARs. Similar enrichment analyses were conducted for gene-proximal and distal (or intergenic) DARs using gene-proximal and distal non-DARs as control.

To compare the TF footprints from different samples, we calculated the count of the ATAC insertions per nucleotide. Then, the number of ATAC-Seq insertions located in a 200-bp window centering at TF motifs was normalized through dividing by the total number of insertions in the flanking 300bp window. For each sample, we averaged the normalized insertions centralizing at TF motif across all footprint sites. To quantitatively compare the footprint, we calculated footprint occupancy score (FOS), following

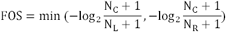

where N_C_ indicates the number of insertions in central region of TF motif. The size of the central region is equal to the length of TF motif. N_L_ and N_R_ are separately 1/3 numbers of the insertions in the left and right flanking regions of TF motif, as an interval 3 times greater than the size of central region was chosen as the size of flanking region.

### Gene expression analysis

Using RNA-Seq, we measured gene expression of retina and RPE from AMD2 left and right eyes. Using Tophat and Cufflinks with default parameters^36^, raw data were mapped to the hg19 genome assembly, and gene expression FPKM (Fragments Per Kilobase Of Exon Per Million) were obtained. In the study, we also used normal, pre-AMD and dry AMD samples from published expression dataset (GEO accession number: GSE29801) ^14^. If multiple ATAC-Seq peaks mapped to the same gene, we select the highest ATAC-Seq peak among the gene-proximal peaks to associate with gene expression. To compare differential chromatin accessibility and differential mRNA expression, we only included genes with the average expression more than 1 FPKM in left and right eyes of AMD2 patient.

### Analysis of GWAS SNPs in DARs

To study the association of DARs with SNPs, we downloaded all SNPs (single nucleotide polymorphisms) from the latest genome-wide association studies (GWAS) performed in AMD^5^. According to genomic location, GWAS SNPs were assigned to corresponding ATAC-Seq peaks. We choose the minimum P value of SNPs in each peak region as the significance of ATAC-Seq peak associated with GWAS SNPs. For ATAC-Seq peak without any SNP, P value of peak was set to 1. Similar analyses were performed for DARs, non-DARs and non-peaks. The flank regions with the same size of peaks were chose as non-peaks. We then calculated the proportion of DARs, non-DARs and non-peaks whose P values are less than a fixed threshold.

### Cell type-specific genes involving in DARs

We collected seven sets of genes which are specifically expressed in rods, cones, horizontal cells, bipolar cells, Müller glia, amacrine cells and ganglion cells, which were assembled from previously published data in mouse^37-44^. Using information downloaded from MGI database (http://www.informatics.jax.org/), we mapped these genes to their corresponding human orthologues. We then compared cell type-specific genes to genes in DARs, and identified the number of cell type-specific genes in associated with DARs.

### Human RPE derived from induced pluripotent stem cells (iPSC) and plasmid transfection

Human induced pluripotent stem cells (iPSC) were cultured and differentiated into RPE monolayers as previously described^45,46^. Briefly, the iPSC line EP1 were maintained on growth factor-reduced Matrigel (BD Biosciences) in mTeSR1 medium (Stem Cell Technologies), in a 10% CO_2_ and 5% O_2_ incubator and amplified by clonal propagation using the ROCK pathway inhibitor Blebbistatin (Sigma). For differentiation, iPSC were plated at higher density (25,000 cells per cm2) and maintained in mTeSR1 to form a monolayer and the culture medium was replaced with differentiation medium (DM) for 40-45 days. Differentiating cells were enzymatically dissociated using 0.25% (wt/vol) collagenase IV (Gibco) and resuspended in AccuMAX (Sigma-Aldrich) to make single cell suspension. Cells were re-plated in fresh Matrigel coated plates and maintained in RPE medium [70% DMEM, 30% Ham’s F12 nutrient mix, 2% B27-serum-free supplement liquid, 1% antibiotic-antimycotic solution (Invitrogen)] for 2-3 months to form mature RPE monolayers.

Immunostaining for RPE-specific markers were performed using the IntraPrep Permeabilization kit (Beckman Coulter) following the manufacturer’s instructions. Primary antibody concentration was 0.046 μg per 1 million cells for mouse anti-RPE65 (Abcam, ab13826) and mouse anti-RLBP1 (Abcam, ab15051). Goat anti-mouse conjugated to Alexa 647 (Invitrogen) was used as a secondary antibody. A nonspecific, species-appropriate isotype control was included in all flow cytometry experiments and stained cells were analyzed using a C6 flow cytometer (Accuri™).

Plasmid transfections were performed using DNA-In Stem (2 uL/well) Transfection Reagent (MTI-Global Stem) in mature iPSC-RPE monolayers (> 2-month-old) cultured in 24 well plates. 1μg of plasmid DNA containing empty vector (pCAGIG) or expression vector pCAGIG-HDAC11 were used for transfection. Plasmid DNA-DNA-In Stem reagent complex is prepared in OptiMEM/RPE medium without antibiotics and incubated for 30 min at RT. Plasmid DNA-DNA-In Stem reagent complex mixture is directly added to the RPE monolayers. Cells were harvested at 72 hours after transfection for total protein extraction.

### Cigarette smoke treatment of iPSC-derived RPE cells

Differentiated RPE cells were plated at 100,000 cells per cm2 on Matrigel-coated plates and allowed to grow for 4 months in RPE medium, consisting of 70% DMEM (Invitrogen), 30% Ham’s F-12 Nutrient Mix (catalog no. 11765; Invitrogen), 1× B27 (Invitrogen), and 1×antibiotic-antimycotic (Invitrogen). When cells were fully differentiated after 4 months, they were exposed to Cigarette Smoke Extract (500ug/ml) (Murty Pharmaceuticals, Lexington, KY) for 2 hours a day for 5 days. After 5 days of treatment, chromatin and protein were extracted separately for ATAC-Seq and HDAC11 assay.

### Preparation of cell lysates and Western blotting analysis

iPSC derived RPE monolayers were rinsed twice with ice-cold PBS and collected by scrapping and lysed with RIPA buffer (20-188, Millipore) supplemented with 1% protease inhibitor cocktail (Sigma-Aldrich). Samples were incubated on ice for 30 minutes and vortexed 2~3 times during the incubation. After that, samples were centrifuged at 13,000 g for 20 minutes. The supernatants were mixed with 4×protein sample buffer (Invitrogen) plus 5% 2-mercaptoethanol (Sigma-Aldrich) and heated at 100°C for 10 minutes. For Western blotting, the procedure was described previously^47^. Briefly, each sample was loaded onto a 4-12% Bis-Tris Nu-PAGE gel (Invitrogen) and transferred to nitrocellulose membranes. Blots were incubated with anti-HDAC11 antibody (1:1000, Abcam, ab18973) or β-tubulin antibody (1:1000, Cell signaling, #2128) overnight at 4°C followed by incubation with HRP-conjugated secondary antibodies (Kirkegaard and Perry Laboratories) for 1 hour at room temperature. Blots were visualized using the Amersham ECL Prime Western Blotting Detection Reagent (GE healthcare life sciences).

Coomassie staining was performed to detect total protein of each sample. After electrophoresis, prefix gel in 50% methanol and 7% acetic acid solution for 15 minutes. After washing, gels were incubated in GelCode Blue Stain Reagent (Thermo Fisher Scientific™) for 1 hour and then destained by washing in water with several changes over 1-2 hours. Gels were scanned and the densitometric analysis was done by Quantity One software (Bio-Rad Laboratories).

### Statistical analysis

All statistical analyses were performed in R platform (https://www.R-project.org/). Fisher’s exact test, Student’s t-test, one-way ANOVA, Pearson’s correlation coefficient were used to assess the significance. Fold change, P-values and FDR (false discovery rate) were calculated in analysis.

## Data availability

The data of ATAC-Seq and RNA-Seq data in this study have been deposited in NCBI’s Gene Expression Omnibus (GEO) under accession number GSE99287.

## Supplementary Information

is available in the online version of the paper.

## Acknowledgements

This work was supported by NIH grants R01EY020560 (to S.B.), R01EY024580 (to J.Q.), R01EY023188 (to S.L.M. and J.Q.), R01EY027691 and Macular Degeneration Foundation (to J.T.H.). We thank Akrit Sodhi, Jeff Mumm, and Albert Jun for insightful discussions.

## Author Contributions

J.Q. and S.B. proposed the project. J.W., C.Z. and H.J. conceived of the method. C.Z., P.S., S.R.S, P. Z. and M.C. conducted experiments. J.W. performed data analysis. J.Q., S.B., J.W. and C.Z. wrote the manuscript. J.Q., S.B., J.W., C.Z., H.J., S.L.M., D.J.Z., J.T.H. and D.S. revised the manuscript. J.Q. and S.B. supervised this work.

## Author Information

All ATAC-Seq and RNA-Seq data has been deposited in GEO under the accession number GSE99287. Correspondence and requests for materials should be addressed to S.B. or J.Q.

## References

1. Jager, R.D., Mieler, W.F. & Miller, J.W. Age-related macular degeneration. N Engl J Med 358, 2606–17 (2008).

2. Wong, W.L. et al. Global prevalence of age-related macular degeneration and disease burden projection for 2020 and 2040: a systematic review and meta-analysis. The Lancet Global Health 2, e106–e116 (2014).

3. Lim, L.S., Mitchell, P., Seddon, J.M., Holz, F.G. & Wong, T.Y. Age-related macular degeneration. Lancet 379, 1728–38 (2012).

4. Ambati, J. & Fowler, B.J. Mechanisms of age-related macular degeneration. Neuron 75, 26–39 (2012).

5. Fritsche, L.G. et al. A large genome-wide association study of age-related macular degeneration highlights contributions of rare and common variants. Nat Genet 48, 134–43 (2016).

6. Fritsche, L.G. et al. Seven new loci associated with age-related macular degeneration. Nat Genet 45, 433–9, 439e1–2 (2013).

7. Seddon, J.M., Reynolds, R. & Rosner, B. Associations of smoking, body mass index, dietary lutein, and the LIPC gene variant rs10468017 with advanced age-related macular degeneration. Mol Vis 16, 2412–24 (2010).

8. Seddon, J.M., George, S., Rosner, B. & Klein, M.L. CFH gene variant, Y402H, and smoking, body mass index, environmental associations with advanced age-related macular degeneration. Hum Hered 61, 157–65 (2006).

9. Buenrostro, J.D., Giresi, P.G., Zaba, L.C., Chang, H.Y. & Greenleaf, W.J. Transposition of native chromatin for fast and sensitive epigenomic profiling of open chromatin, DNA-binding proteins and nucleosome position. Nat Methods 10, 1213–8 (2013).

10. Bhutto, I. & Lutty, G. Understanding age-related macular degeneration (AMD): relationships between the photoreceptor/retinal pigment epithelium/Bruch’s membrane/choriocapillaris complex. Mol Aspects Med 33, 295–317 (2012).

11. Freund, C.L. et al. De novo mutations in the CRX homeobox gene associated with Leber congenital amaurosis. Nat Genet 18, 311–2 (1998).

12. Nishida, A. et al. Otx2 homeobox gene controls retinal photoreceptor cell fate and pineal gland development. Nat Neurosci 6, 1255–63 (2003).

13. Woodell, A. & Rohrer, B. A mechanistic review of cigarette smoke and age-related macular degeneration. Adv Exp Med Biol 801, 301–7 (2014).

14. Newman, A.M. et al. Systems-level analysis of age-related macular degeneration reveals global biomarkers and phenotype-specific functional networks. Genome Medicine 4, 1 (2012).

15. Denny, S.K. et al. Nfib Promotes Metastasis through a Widespread Increase in Chromatin Accessibility. Cell 166, 328–42 (2016).

16. Koppenol, W.H., Bounds, P.L. & Dang, C.V. Otto Warburg’s contributions to current concepts of cancer metabolism. Nat Rev Cancer 11, 325–37 (2011).

17. Levine, A.J. & Puzio-Kuter, A.M. The control of the metabolic switch in cancers by oncogenes and tumor suppressor genes. Science 330, 1340–4 (2010).

18. Su, B. et al. Abnormal mitochondrial dynamics and neurodegenerative diseases. Biochim Biophys Acta 1802, 135–42 (2010).

19. Lin, M.T. & Beal, M.F. Mitochondrial dysfunction and oxidative stress in neurodegenerative diseases. Nature 443, 787–95 (2006).

20. Fan, J., Krautkramer, K.A., Feldman, J.L. & Denu, J.M. Metabolic regulation of histone post-translational modifications. ACS Chem Biol 10, 95–108 (2015).

21. Bolger, A.M., Lohse, M. & Usadel, B. Trimmomatic: a flexible trimmer for Illumina sequence data. Bioinformatics, btu170 (2014).

22. Langmead, B. & Salzberg, S.L. Fast gapped-read alignment with Bowtie 2. Nature methods 9, 357–359 (2012).

23. Li, H. et al. The sequence alignment/map format and SAMtools. Bioinformatics 25, 2078–2079 (2009).

24. Zhang, Y. et al. Model-based analysis of ChIP-Seq (MACS). Genome biology 9, 1 (2008).

25. Anders, S., Pyl, P.T. & Huber, W. HTSeq–a Python framework to work with high-throughput sequencing data. Bioinformatics, btu638 (2014).

26. Krzywinski, M. et al. Circos: an information aesthetic for comparative genomics. Genome research 19, 1639–1645 (2009).

27. Thurman, R.E. et al. The accessible chromatin landscape of the human genome. Nature 489, 75–82 (2012).

28. Robinson, M.D., McCarthy, D.J. & Smyth, G.K. edgeR: a Bioconductor package for differential expression analysis of digital gene expression data. Bioinformatics 26, 139–140 (2010).

29. Wang, K., Li, M. & Hakonarson, H. ANNOVAR: functional annotation of genetic variants from high-throughput sequencing data. Nucleic acids research 38, e164– e164 (2010).

30. Subramanian, A. et al. Gene set enrichment analysis: a knowledge-based approach for interpreting genome-wide expression profiles. Proceedings of the National Academy of Sciences of the United States of America 102, 15545–15550 (2005).

31. Wright, A.F., Chakarova, C.F., Abd El-Aziz, M.M. & Bhattacharya, S.S. Photoreceptor degeneration: genetic and mechanistic dissection of a complex trait. Nat Rev Genet 11, 273–84 (2010).

32. Strunnikova, N.V. et al. Transcriptome analysis and molecular signature of human retinal pigment epithelium. Hum Mol Genet 19, 2468–86 (2010).

33. Matys, V. et al. TRANSFAC: transcriptional regulation, from patterns to profiles. Nucleic Acids Res 31, 374–8 (2003).

34. Grant, C.E., Bailey, T.L. & Noble, W.S. FIMO: scanning for occurrences of a given motif. Bioinformatics 27, 1017–8 (2011).

35. Sung, M.H., Guertin, M.J., Baek, S. & Hager, G.L. DNase footprint signatures are dictated by factor dynamics and DNA sequence. Mol Cell 56, 275–85 (2014).

36. Trapnell, C. et al. Differential gene and transcript expression analysis of RNA-seq experiments with TopHat and Cufflinks. Nature protocols 7, 562–578 (2012).

37. Siegert, S. et al. Transcriptional code and disease map for adult retinal cell types. Nat Neurosci 15, 487–95, S1–2 (2012).

38. Macosko, E.Z. et al. Highly Parallel Genome-wide Expression Profiling of Individual Cells Using Nanoliter Droplets. Cell 161, 1202–14 (2015).

39. Hsiau, T.H. et al. The cis-regulatory logic of the mammalian photoreceptor transcriptional network. PLoS One 2, e643 (2007).

40. Trimarchi, J.M. et al. Molecular heterogeneity of developing retinal ganglion and amacrine cells revealed through single cell gene expression profiling. J Comp Neurol 502, 1047–65 (2007).

41. Roesch, K. et al. The transcriptome of retinal Muller glial cells. J Comp Neurol 509, 225–38 (2008).

42. Hao, H. et al. Transcriptional regulation of rod photoreceptor homeostasis revealed by in vivo NRL targetome analysis. PLoS Genet 8, e1002649 (2012).

43. Blackshaw, S., Fraioli, R.E., Furukawa, T. & Cepko, C.L. Comprehensive analysis of photoreceptor gene expression and the identification of candidate retinal disease genes. Cell 107, 579–89 (2001).

44. Blackshaw, S. et al. Genomic analysis of mouse retinal development. PLoS Biol 2, E247 (2004).

45. Maruotti, J. et al. A simple and scalable process for the differentiation of retinal pigment epithelium from human pluripotent stem cells. Stem Cells Transl Med 2, 341–54 (2013).

46. Maruotti, J. et al. Small-molecule-directed, efficient generation of retinal pigment epithelium from human pluripotent stem cells. Proc Natl Acad Sci U S A 112, 10950–5 (2015).

47. Shang, P. et al. The amino acid transporter SLC36A4 regulates the amino acid pool in retinal pigmented epithelial cells and mediates the mechanistic target of rapamycin, complex 1 signaling. Aging Cell 16, 349–359 (2017).

