## Supplementary Materials for "A widespread decrease of chromatin accessibility in age-related macular degeneration"

### Extended Data Tables

**Extended Data Table 1.** The characteristics of donors

| Variable | Norm (n = 5) | AMD (n = 5) | P value* |
|---|---|---|---|
| Gender (male) | 2 (40%) | 1 (20%) | 0.41 |
| Age (years) | 84: 78 ~ 92 | 90: 85 ~ 95.5 | 0.24 |
| No. of eyes | 8 | 8 | 0.99 |
| Interval (hours)& | 8.6 | 6 | 0.24 |

*fisher's exact test and Student's t-test were performed respectively. &Interval indicates the time from death to procurement of eye.

**Extended Data Table 2.** Statistics of ATAC-Seq for retina and RPE samples

| Sample* | Properly paired fragments (%)# | Qualified fragments& | Sample* | Properly paired fragments (%)# | Qualified fragments& |
|---|---|---|---|---|---|
| NOR1_Retina_MacR | 95,198,063 (77.6) | 56,151,841 | AMD5_Retina_MacR | 36,530,166 (86.8) | 22,515,903 |
| NOR1_Retina_PerL | 30,375,713 (84.0) | 23,357,162 | AMD5_Retina_PerR | 61,929,860 (83.5) | 47,717,112 |
| NOR1_Retina_PerR | 66,360,580 (81.7) | 41,210,030 | NOR2_RPE_Mac | 62,240,653 (86.6) | 38,332,398 |
| NOR2_Retina_Mac | 47,063,121 (82.8) | 31,460,883 | NOR2_RPE_PerL | 83,058,819 (83.6) | 44,954,984 |
| NOR2_Retina_Per | 61,502,065 (85.6) | 46,540,872 | NOR2_RPE_PerR | 54,504,714 (73.5) | 30,156,496 |
| NOR3_Retina_MacR | 51,261,953 (84.2) | 32,235,358 | NOR3_RPE_PerR | 47,401,144 (83.3) | 35,286,709 |
| NOR4_Retina_MacL | 73,794,675 (77.9) | 33,234,136 | NOR4_RPE_MacL | 80,538,584 (70.6) | 55,001,840 |
| NOR4_Retina_MacR | 56,130,375 (79.4) | 24,316,121 | NOR4_RPE_MacR | 87,406,027 (73.3) | 47,954,810 |
| NOR4_Retina_PerL | 44,670,935 (79.4) | 26,770,972 | NOR4_RPE_PerL | 95,699,676 (69.6) | 65,998,781 |
| NOR4_Retina_PerR | 54,900,814 (76.3) | 32,758,426 | NOR4_RPE_PerR | 95,508,108 (68.1) | 60,201,309 |
| NOR5_Retina_PerR | 71,799,205 (90.5) | 45,054,235 | AMD1_RPE_MacL | 46,657,027 (85.3) | 31,307,075 |
| AMD1_Retina_MacL | 53,298,282 (81.3) | 36,206,550 | AMD1_RPE_MacR | 63,649,499 (81.0) | 36,630,371 |
| AMD1_Retina_MacR | 42,169,255 (82.0) | 26,416,356 | AMD1_RPE_PerR | 57,068,660 (80.9) | 38,720,478 |
| AMD1_Retina_PerR | 59,425,134 (85.1) | 38,986,244 | AMD2_RPE_MacL | 80,189,619 (75.9) | 41,553,090 |
| AMD2_Retina_MacL | 52,973,396 (76.4) | 25,863,986 | AMD2_RPE_PerL | 64,525,397 (67.8) | 35,411,272 |
| AMD2_Retina_MacR | 41,334,371 (75.6) | 20,019,283 | AMD2_RPE_PerR | 65,795,447 (65.1) | 39,301,200 |
| AMD2_Retina_PerL | 41,470,433 (77.4) | 23,218,933 | AMD3_RPE_MacR | 72,938,776 (78.6) | 31,369,459 |
| AMD2_Retina_PerR | 72,157,524 (69.2) | 24,251,972 | AMD3_RPE_PerR | 68,690,135 (75.0) | 32,083,977 |
| AMD3_Retina_PerR | 116,652,520 (86.1) | 38,580,810 | AMD4_RPE_MacR | 32,335,897 (83.3) | 20,123,768 |

| | | | | | |
|---|---|---|---|---|---|
| AMD4_Retina_MacR | 39,004,966 (83.0) | 19,575,730 | AMD5_RPE_MacL | 57,902,954 (81.4) | 36,203,385 |
| AMD4_Retina_PerR_ Rep1 | 51,899,554 (78.4) | 38,903,055 | AMD5_RPE_MacR | 42,040,435 (82.3) | 32,761,740 |
| AMD4_Retina_PerR Rep2 | 48,746,337 (79.7) | 34,160,635 | AMD5_RPE_PerR | 48,591,576 (82.6) | 48,591,576 |
| AMD5_Retina_MacL | 55,453,128 (85.5) | 35,550,123 | | | |
| AMD5_Retina_PerL | 45,593,306 (86.1) | 32,117,341 | Average | 60,400,845 (78.5) | 35,799,881 |

[*]The name of sample is consisted of disease status of donors (NOR: normal; AMD: AMD), type of tissue (Retina vs RPE), region (Mac: macula; Per: periphery) and eye index (R: right eye; L: left eye). For the sample of AMD4_Retina_PerR, there are two replicates (Rep1 and Rep2). [#]include fragments from chromosome M and Y. The percentage in the bracket represents the ratio of properly paired fragments in total raw fragments. [&]the quantified fragments are those fragments of MAPQ score > 10 after removing fragments from chromosome M and Y as well as duplicate fragments.

**Extended Data Table 3.** The enriched TF motifs in footprints within DARs of retinas and RPE

| Transcription factor | Motif ID | FDR | | | Transcription factor | Motif ID | FDR | | |
|---|---|---|---|---|---|---|---|---|---|
| | | All | Proximal | Distal | | | All | Proximal | Distal |
| **From footprints within DARs of retinas** | | | | | | | | | |
| OTX2 | M01719 | 7E-14 | 3E-3 | 3E-13 | CRX | M00623 | 1E-6 | 0.04 | 1E-6 |
| POU2F1 | M00138 | 2E-6 | 0.09 | 4E-7 | MEF2D | M00941 | 1E-4 | 8E-3 | 0.07 |
| PAX6 | M01391 | 5E-4 | 0.57 | 6E-6 | FOXJ2 | M00422 | 7E-4 | 8E-3 | 0.34 |
| POU3F3 | M01324 | 1E-3 | 0.09 | 0.01 | TLX2 | M01420 | 2E-3 | 0.23 | 3E-3 |
| CDC5L | M00478 | 4E-3 | 0.01 | 0.63 | RORA | M00157 | 8E-3 | 0.14 | 0.12 |
| CUX1 | M01344 | 9E-3 | 6E-3 | 0.99 | RXRA | M01152 | 0.01 | 0.04 | 0.53 |
| FOXO3 | M01137 | 0.01 | 0.09 | 0.37 | VAX2 | M01327 | 0.01 | 0.57 | 2E-3 |
| NKX3-1 | M00451 | 0.01 | 0.03 | 0.68 | PHOX2A | M01444 | 0.01 | 0.53 | 2E-3 |
| MEIS1 | M00420 | 0.02 | 0.09 | 0.51 | POU6F1 | M00465 | 0.02 | 0.32 | 0.06 |
| ONECUT1, ONECUT2 | M00639 | 0.03 | 0.25 | 0.11 | IRX5 | M01472 | 0.03 | 0.15 | 0.34 |
| ZBTB16 | M01075 | 0.04 | 0.08 | 0.90 | GTF2IRD1 | M01229 | 0.05 | 0.24 | 0.25 |
| **From footprints within DARs of RPE** | | | | | | | | | |
| POU3F3 | M01324 | 2E-4 | 0.20 | 8E-8 | SOX5 | M00042 | 2E-3 | 0.38 | 3E-9 |
| SOX4 | M01308 | 2E-3 | 0.02 | 0.07 | CTNNB1 | M03539 | 0.01 | 0.01 | 0.90 |
| POU2F1 | M00161 | 0.02 | 0.62 | 3E-6 | PITX2 | M01447 | 0.02 | 0.52 | 5E-5 |
| CUX1 | M00104 | 0.02 | 6E-4 | 0.99 | NKX3-1 | M00451 | 0.02 | 0.09 | 0.11 |
| CEBPG | M00622 | 0.02 | 0.06 | 0.27 | OTX2 | M01719 | 0.02 | 0.46 | 3E-4 |

| | | | | | | | | | |
|---|---|---|---|---|---|---|---|---|---|
| SATB1 | M01723 | 0.02 | 0.64 | 1E-5 | CDC5L | M00478 | 0.04 | 0.61 | 3E-3 |
| TCF4 | M02033 | 0.04 | 0.13 | 0.21 | | | | | |

Note, According to the genomic location of peaks, all peaks were divided into two groups including those proximal and distal to gene body regions. We calculated the FDR for proximal and distal peaks, respectively.

**Extended Data Table 4.** The expression of histone deacetylases in RPE tissue

| Gene | Probe ID | Normal (n = 96) | AMD (n = 30) | Log2 fold change | P value |
|---|---|---|---|---|---|
| HDAC1 | 12245 | 8.56 | 8.53 | -0.04 | 0.54 |
| HDAC2 | 40294 | 8.67 | 8.59 | -0.08 | 0.30 |
| HDAC3 | 29325 | 3.90 | 3.94 | 0.04 | 0.61 |
| HDAC4 | 14394 | 5.49 | 5.61 | 0.12 | 0.20 |
| HDAC5 | 3947 | 7.66 | 7.55 | -0.11 | 0.23 |
| HDAC6 | 25939 | 7.59 | 7.53 | -0.06 | 0.39 |
| HDAC7 | 9457 | 9.34 | 9.33 | 0.00 | 0.97 |
| HDAC8 | 24646 | 4.15 | 4.20 | 0.04 | 0.61 |
| HDAC9 | 33494 | 2.52 | 2.61 | 0.09 | 0.45 |
| HDAC10 | 19616 | 3.53 | 3.24 | -0.29 | **0.02** |
| HDAC11 | 38006 | 4.17 | 4.41 | 0.24 | **0.05** |
| SIRT1 | 41246 | 5.34 | 5.10 | -0.24 | **0.02** |
| SIRT2 | 5854 | 6.08 | 5.99 | -0.09 | 0.24 |
| SIRT3 | 8468 | 7.32 | 7.46 | 0.14 | 0.13 |
| SIRT4 | 43993 | 3.13 | 3.13 | -0.01 | 0.97 |
| SIRT5 | 18987 | 6.31 | 6.18 | -0.13 | 0.14 |
| SIRT6 | 11898 | 5.17 | 5.19 | 0.02 | 0.76 |
| SIRT7 | 26965 | 6.66 | 6.55 | -0.11 | 0.25 |

Note, gene expression was obtained from the microarray GSE29801. RPE samples from both macular and peripheral regions were included. Here AMD samples indicate the dry AMD.

**Extended Data Figures**

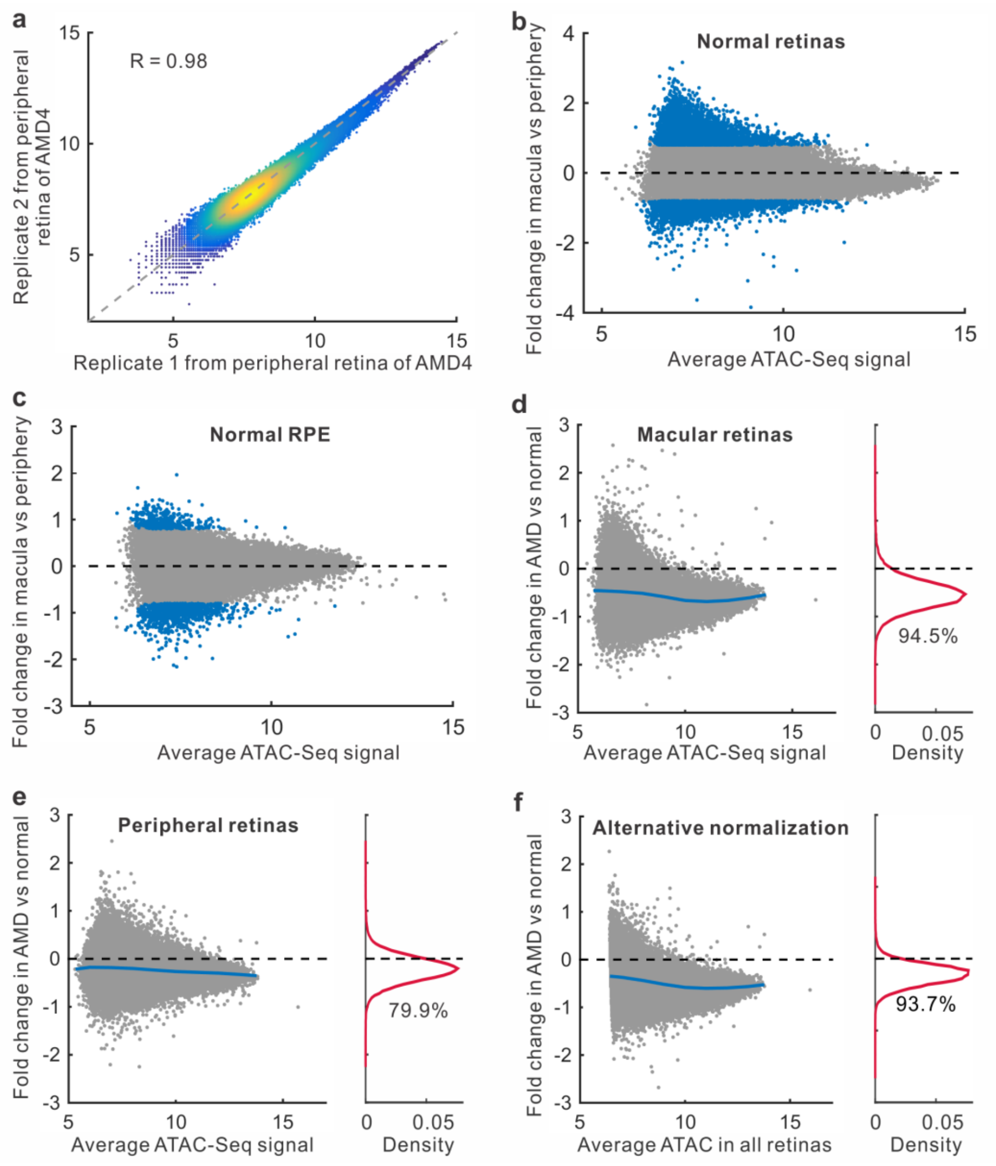


**Extended Data Figure 1.** Region-specific peaks and changes in chromatin accessibility. (a) The replicated samples from adjacent regions in peripheral retina of right eye of one AMD patient. The color represents the density of peaks. (b-c) Changes of chromatin accessibility in macular regions of retinas or RPE from healthy donors. The blue dots are peaks with significantly differential accessibility. (d-e) Changes of chromatin accessibility in AMD relative to normal in retina samples from macular and peripheral regions, respectively. Each dot represents one ATAC-Seq peak. Blue line indicates average fold changes of peaks. The percentage of reduced peaks is shown under the density curve. (f) Changes of chromatin accessibility in all AMD retinas. For alternative normalization, we took the count of the properly paired fragments as the library size.

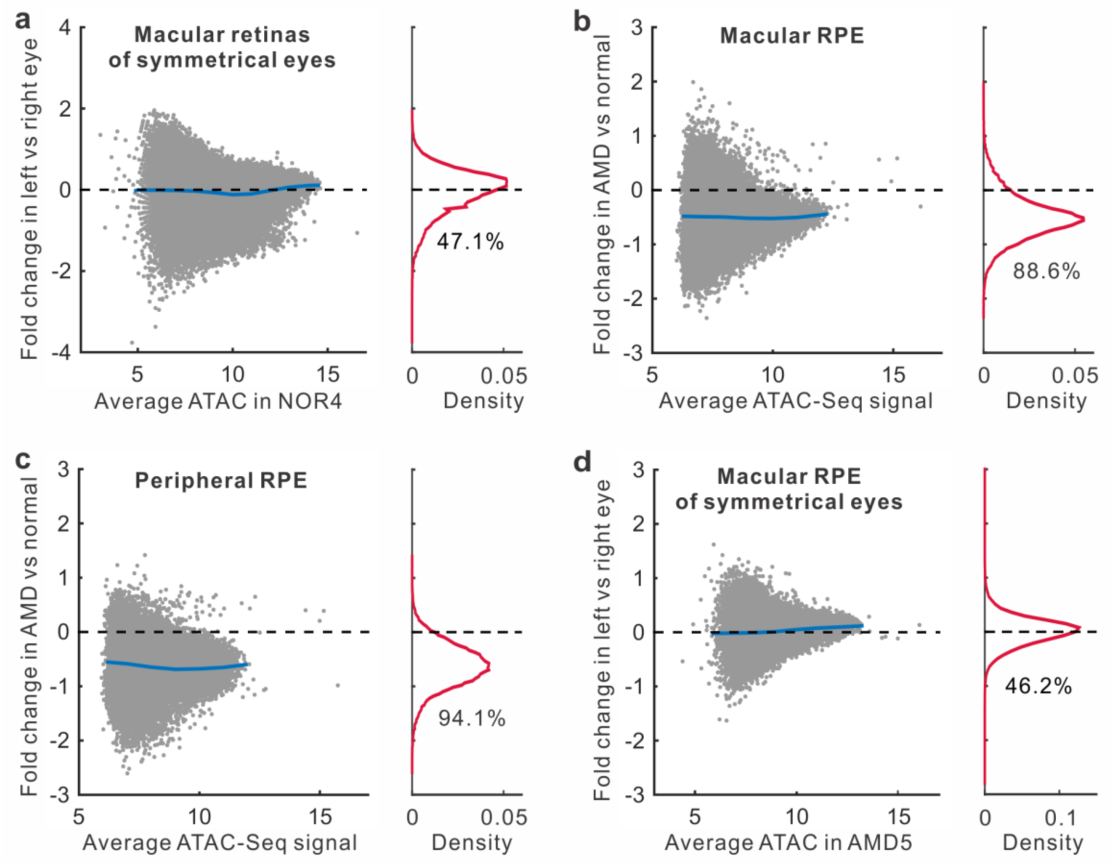


**Extended Data Figure 2.** Changes of chromatin accessibility in eyes from the same donor and region-specific RPE samples. (a) Changes of chromatin accessibility in macular retinas from the donors whose eyes are at the same stage of disease. The blue line represents the average change of ATAC-Seq peaks (grey dots). The percentage of peaks with the reduced accessibility is showed under the density curve. (b-c) Changes of chromatin accessibility in AMD relative to normal in RPE samples from macular and peripheral regions, respectively. (d) Changes of chromatin accessibility in macular RPE from the donors whose eyes are at the same stage of disease.

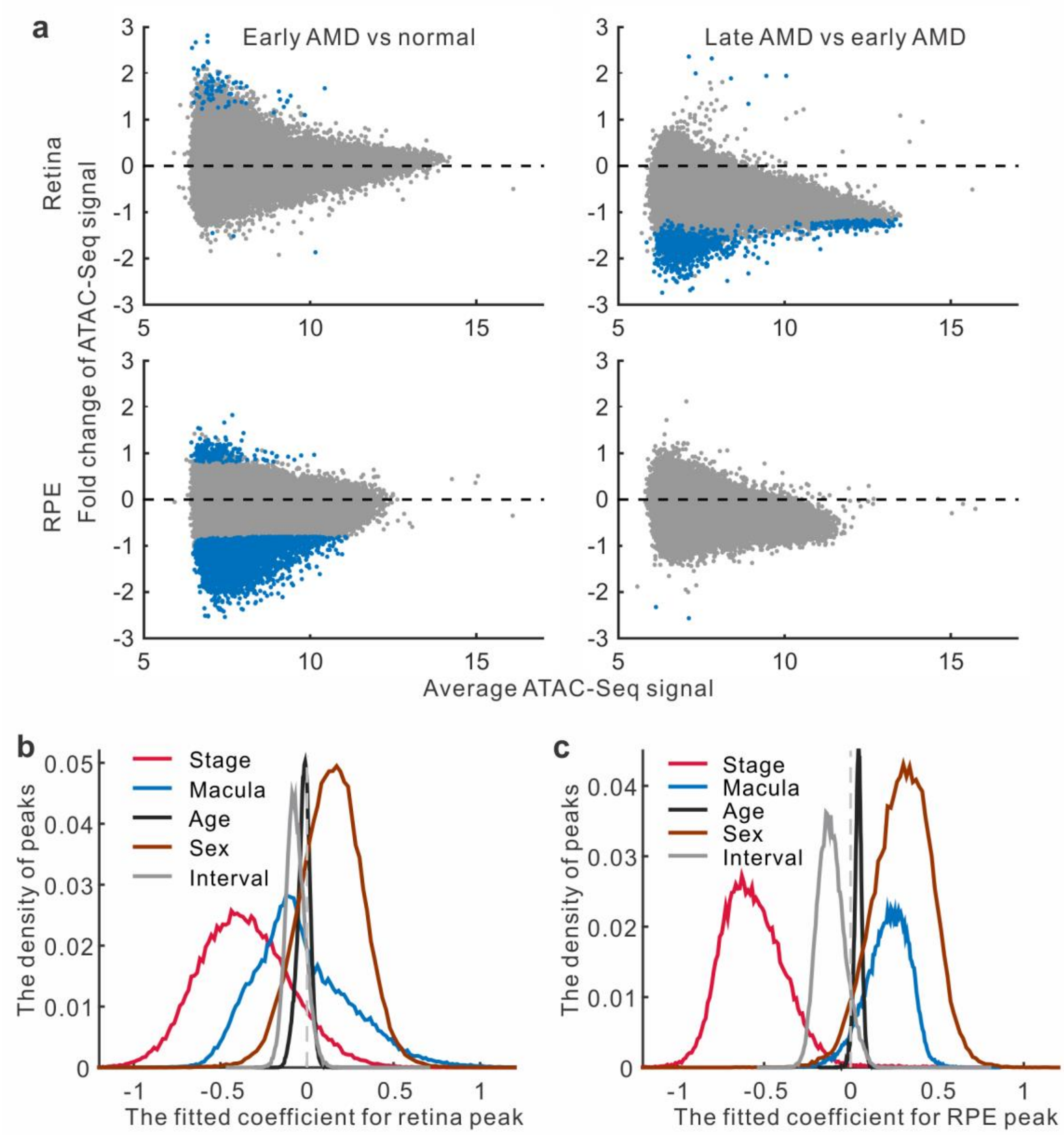


**Extended Data Figure 3.** Significant changes in chromatin accessibility at different stages of disease and models of retina and RPE peaks. (a) Changes of ATAC-Seq signal at different stages of AMD. The blue dots are peaks with significantly differential chromatin accessibility. (b-c) The density of the coefficients in the fitting model of peaks from retina and RPE samples. For each peak, the coefficient of every variable was derived from linear regression model.

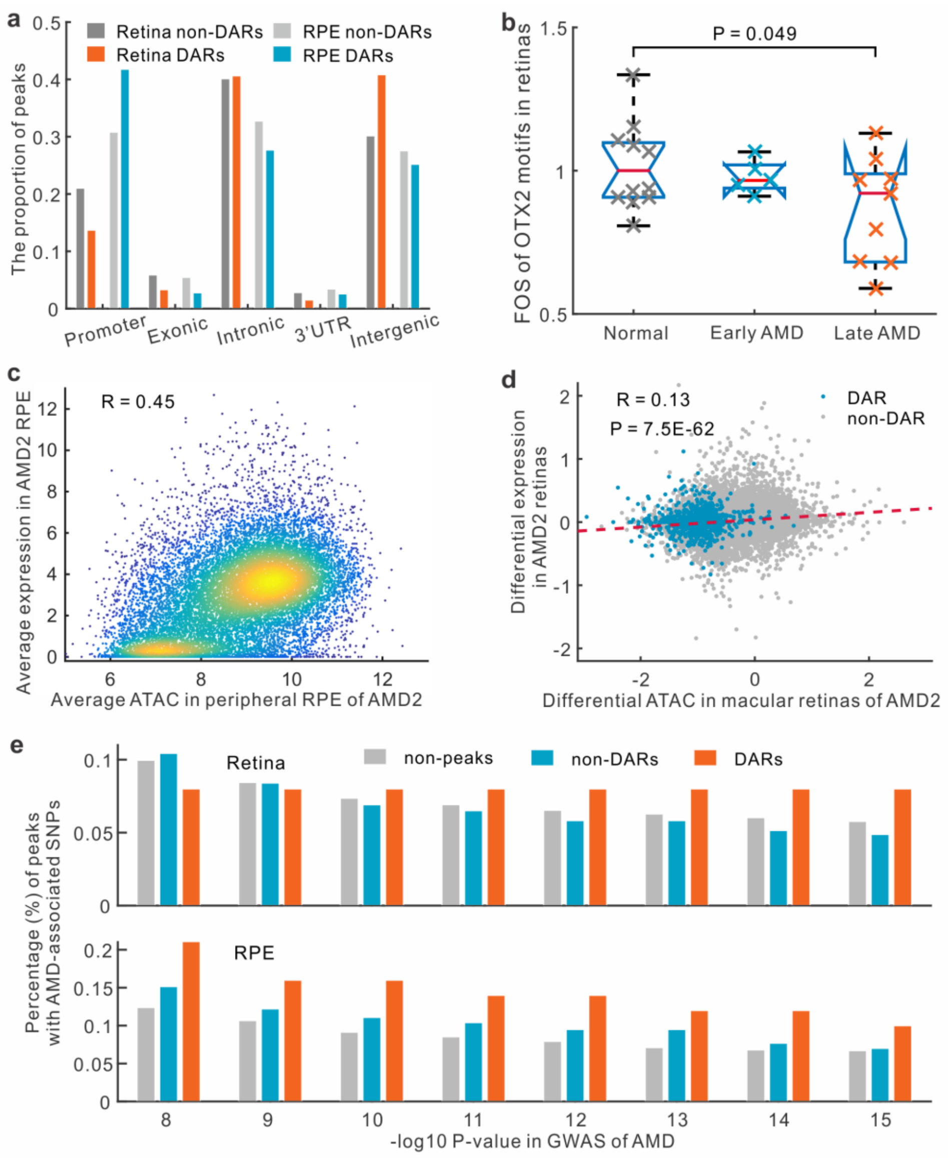


**Extended Data Figure 4.** Features of differentially accessible regions (DARs). (a) The proportion of ATAC-Seq peaks at different genomic locations. 3'UTR, three prime untranslated region. (b) Footprint occupancy scores (FOS) for OTX2 motifs in normal, early-stage, and late-stage AMD retinas. (c) Comparison of chromatin accessibility and gene expression in the RPE of AMD patient. Pearson's correlation coefficient (R) is shown. (d) Comparison of differential accessibility and differential expression in retinas of AMD patient. Comparison of left eye (late-stage AMD) with right eye (early-stage AMD) of the AMD patient was performed for both differential accessibility and differential expression. (e) The percentage of ATAC-Seq peaks with significant GWAS SNPs. SNPs were significant if p value of SNP in GWAS (genome-wide association study) less than a fixed threshold (defined by x-axis).

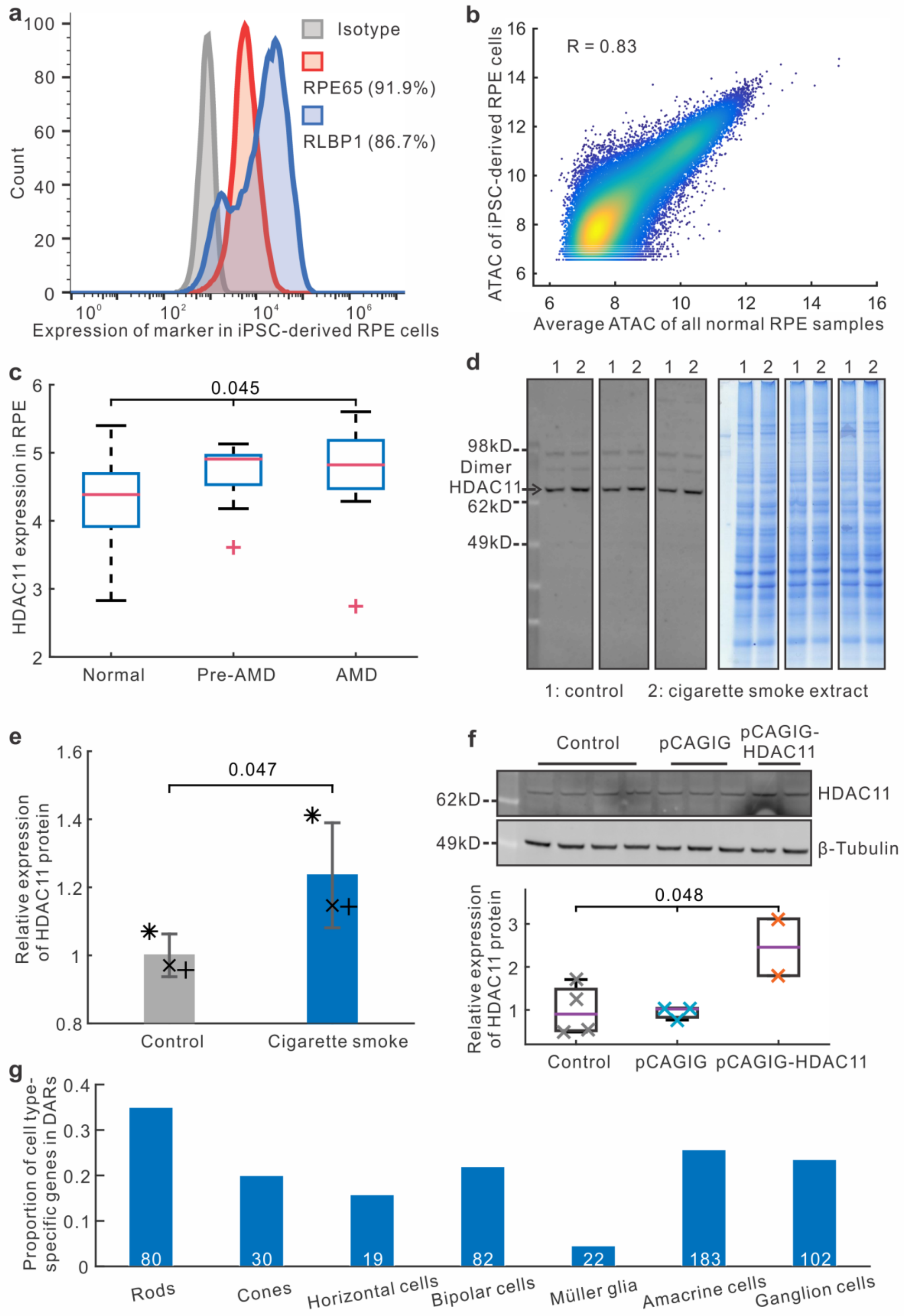
a
Count
Expression of marker in iPSC-derived RPE cells
Isotype
RPE65 (91.9%)
RLBP1 (86.7%)
b
R = 0.83
ATAC of iPSC-derived RPE cells
Average ATAC of all normal RPE samples
c
0.045
HDAC11 expression in RPE
Normal
Pre-AMD
AMD
d
98kD
Dimer
HDAC11
62kD
49kD
1: control
2: cigarette smoke extract
e
0.047
Relative expression of HDAC11 protein
Control
Cigarette smoke
f
Control
pCAGIG
pCAGIG-HDAC11
62kD
HDAC11
49kD
β-Tubulin
0.048
Relative expression of HDAC11 protein
Control
pCAGIG
pCAGIG-HDAC11
g
Proportion of cell type-specific genes in DARs
80
30
19
82
22
183
102
Rods
Cones
Horizontal cells
Bipolar cells
Müller glia
Amacrine cells
Ganglion cells

**Extended Data Figure 5.** HDAC11 change in iPSC-derived RPE cells. (a) Flow cytometric analysis of the expression of RPE specific markers RPE65 and RLBP1 from 12-week-old iPSC-derived RPE monolayers. (b) Comparison of chromatin accessibility in RPE tissue of healthy donors and human iPSC-derived RPE cells. (c) *HDAC11* expression in peripheral RPE at different disease stages. The data are from microarray GSE29801. (d) Western blot of HDAC11 under cigarette smoke treatment in human iPSC-derived RPE cells. (e) Statistics on HDAC11 expression under cigarette smoke treatment. Paired t-test was performed. (f) Abundance of HDAC11 in control, empty vector (pCAGIG), and HDAC11 overexpression (pCAGIG-HDAC11). Top panel, Western blot. Bottom panel, statistics of Western blot and one-way ANOVA was performed. (g) The proportion of cell type-specific genes located in retinal differentially accessible regions (DARs). The number of cell type-specific genes in the study was shown at the bottom of the bar.
